# Stability of microbiota in vineyard soils across consecutive years

**DOI:** 10.1101/2021.04.29.442071

**Authors:** Alessandro Cestaro, Emanuela Coller, Davide Albanese, Erika Stefani, Massimo Pindo, Cluadio Ioriatti, Roberto Zanzotti, Caludio Donati

## Abstract

Agricultural soils harbor rich and diverse microbial communities that have a deep influence on soil properties and productivity. Large scale studies have shown the impact of environmental parameters like climate or chemical composition on the distribution of bacterial and fungal species. Comparatively, little data exists documenting how soil microbial communities change between different years. Quantifying the temporal stability of soil microbial communities will allow us to better understand the relevance of the differences between environments and their impact on ecological processes on the global and local scale.

We characterized the bacterial and fungal components of the soil microbiota in ten vineyards in two consecutive years. Despite differences of species richness and diversity between the two years, we found a general stability of the taxonomic structure of the soil microbiota. Temporal differences were smaller than differences due to geographical location, vineyard land management or differences between sampling sites within the same vineyard. Using machine learning, we demonstrated that each site was characterized by a distinctive microbiota, and we identified a reduced set of indicator species that could classify samples according to their geographic origin across different years with high accuracy.

**Importance:** The temporal stability of the soil microbiota is important to understand the relevance of the differences that are found in response to a variety of environmental factors. By comparing fungal and bacterial microbiota from samples collected in the same sites in two consecutive years, we found a remarkable stability of both components, with characteristic differences between bacteria and fungi. Our work fills an important gap toward the definition of a microbial cartography of agricultural soils.

## Introduction

Soils are colonized by complex and still poorly characterized microbial communities, that constitute a large fraction of the Earth biomass [1], [2]. Thanks to recent technological advancements, cultivation-free approaches have provided a first picture of the diversity of the soil microbiota, although significant knowledge gaps still exist [3], [4]. These studies have shown that soil, together with the plant rhizosphere, is host to the most diverse amongst free living and host-associated microbial communities [5].

A large amount of work has been devoted to the understanding of the ecological factors that drive the composition of soil microbial communities and global scale studies have shown that a relatively small number of ubiquitous taxa dominate the bacterial component of the soil microbiota [6], [7]. Studies on the global distribution of fungal communities show significant diversity that correlate with geography, while a small number of generalist fungal taxa are widespread in the global soil fungal communities [8], [9]. Niche differentiation of soil bacterial and fungal communities is associated with a combination of geographical and environmental variables that often influence their diversity in a contrasting way [3].

Metagenomic characterization of soil microbial communities have received special attention in the context of grape cultivation and wine making [10]. The distinct sensory characteristics that distinguish wines from different regions are the product of a large number of physical and biological factors; amongst these, several studies have documented the role of soil microbial communities [11][12] in determining the microbial contribution to the regional differentiation of wine, in an attempt to define a microbial “Terroir” and how this is influenced by climatic and geographic factors, and management [13], [14]. Despite the fact that, in order to define a microbial contribution to regional characteristics of cultures such as wine, the dynamics of soil microbiota over extended periods of time must be limited compared to differences due to other environmental variables, like, *e.g.* location, climate, or soil chemical characteristics, few studies have addressed the stability of soil microbial communities over time [1]. Studies tracking the variations of soil bacterial communities in different seasons within one year have shown that bacterial communities vary more across land uses than time [15,16]. However, despite the community modifications due to the seasonal changes and / or due to the plant growth cycle, few data are available regarding the stability of bacterial and fungal communities colonizing soil over consecutive years and how variations between years compare to the variability induced by land management or geographic location [17][18][19].

In a previous work, we have used a metabarcoding approach to characterize the bacterial and fungal microbiota of vineyard soils in the Italian Trentino region, finding that land use and location concurrently shape the soil microbiota, with characteristic differences between the bacterial and fungal components [20]. Here, we examine in depth the stability of the soil microbiome across two consecutive years. We show that in both bacterial and fungal communities the variability between the two years is significantly smaller than that related to locations and land use. The majority of bacterial species in each site was present in both years, that differ mainly in the presence/absence of rare taxa. For fungi, we found that in each site a large fraction of taxa with high abundance were present only in one of the two sampling years. Using machine learning we found that a predictor trained on the distribution of taxa in 2017 could classify 2018 samples according to their origin both for bacteria and fungi, showing that both components of the microbiota are highly associated with the geographical origin of the soil.

## Results

After preprocessing and filtering we obtained 10,924,706 16S V4 sequences and 14,672,492 ITS1 sequences, respectively, that were denoised into 30,869 SVs for bacteria and 14,099 SVs for fungi. The samples were rarefied to 15,000 reads per sample, both for Bacteria and Fungi, obtaining a dataset of 5,220,000 reads and 30,815 SVs from 348 samples for 16S and 5,115,000 reads and 14,021 SVs from 341 samples for ITS.

We compared the relative abundances of SVs across samples collected in the same sampling sites in 10 vineyards from 4 different locations situated in the Adige valley in the northern Italian region Trentino [20]. In each vineyard, 6 soil samples were collected between May 4th and May 18th, 2017, and between May 7th and May 24th, 2018, in 3 sampling points, namely between the rows (V) and in the perennial crop area at a distance of 8 (P1) and 16 (P2) meters from the border of the vineyard in each of the two consecutive sampling years. Comparing the meteorological data from 4 meteorological stations located in close proximity of the vineyards, we found that 2018 was characterized by a higher amount of accumulated rain than 2017 (Supplementary Figure S2).

### Diversity and Richness of soil microbiota in two consecutive years

Although most taxa had significant shifts in their relative abundance, we found a general stability of the overall structure of the bacterial microbiota at high taxonomic levels (Figure 1a). In both years the most abundant Phylum was Acidobacteria, followed by Proteobacteria in 2017 and Actinobacteria in 2018. The different proportions of Proteobacteria and Actinobacteria was the only major difference between the two consecutive years at the Phylum level. At the Family level, the most abundant in both years were *Gp6*, followed by *Nitrosospheraceae* and *Planctomycetaceae* (Supplementary Figure S3a). The stability of the bacterial component of the microbiota between the two years was evident by comparing the relative abundance of each Genus separately in each site (Fig. 1b). Confirming the results at the Phylum level, Genuses of the Phyla *Actinobacteria* and *Thaumarcheota* clustered above the diagonal, while Genuses of the Phylum *Proteobacteria* were more numerous below the diagonal. For fungi (Figure 1c), we found that in both years the dominant component of the microbiome was represented by *Ascomycota*, that accounted for more than 50% of the soil mycobiota, followed by *Zygomycota* and *Basidiomycota*. At the Family level, the most abundant in both years were *Mortierellaceae* followed by *Nectriaceae* and a family of unidentified *Ascomycota* (Supplementary Figure S3b). However, the degree of correlation between 2017 and 2018 relative abundances at the Genus level was lower than in Bacteria, as shown by the more dispersed distribution in Fig. 1d.

**Figure 1.**
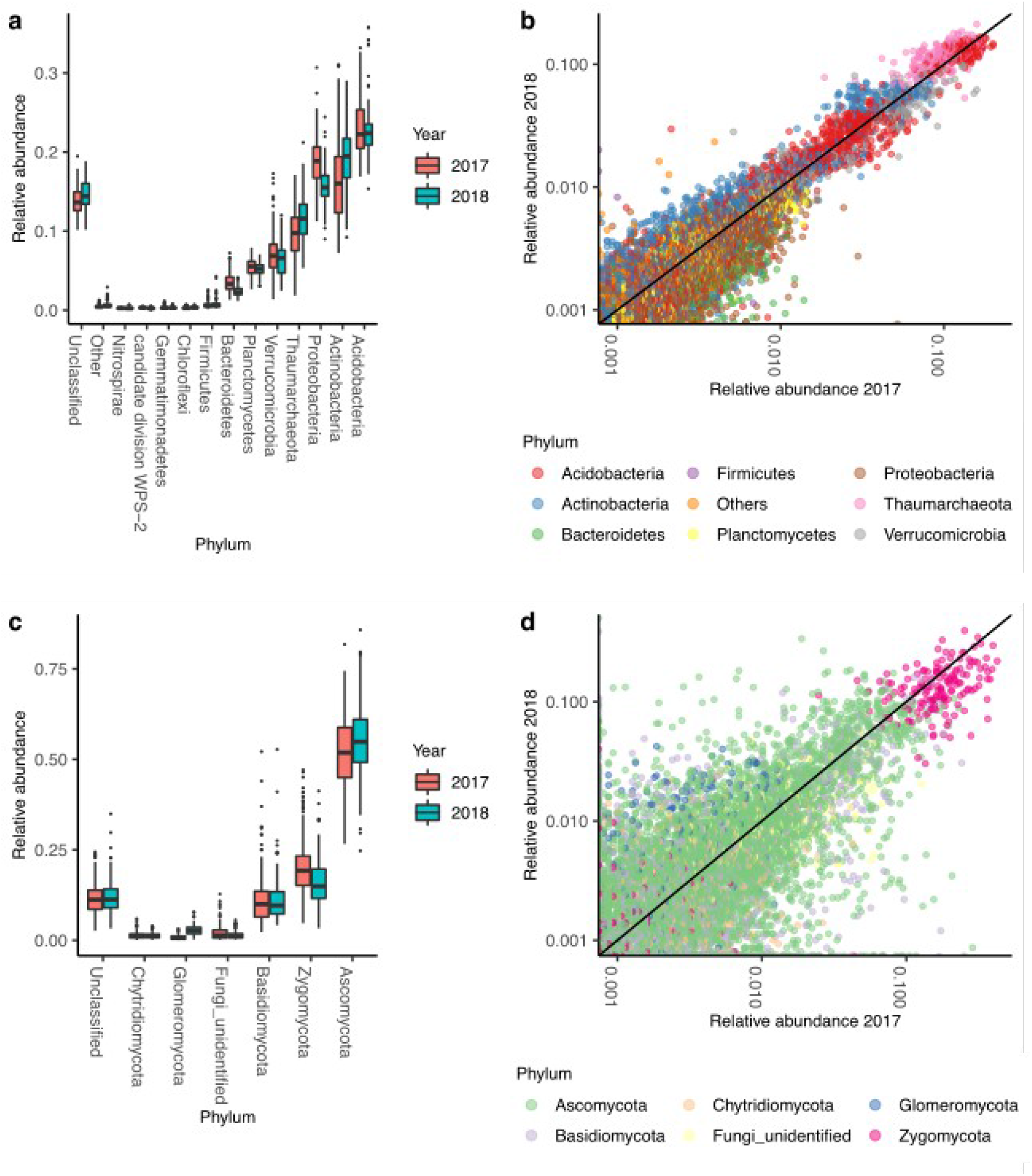
a) Relative abundance of most abundant bacterial phyla in two consecutive years. b) scatter plot of the relative abundances of bacterial SVs in 2018 vs 2017. Each dot represents a bacterial genus in a given site, and horizontal and vertical coordinates are the relative abundances in 2017 and 2018, respectively. The straight line is the bisectrix of the first quadrant. c) same as a), for fungi. d) same as b, for fungi.

To characterize the changes in the structure of the soil microbiome between the two years, we compared the diversity and richness of the microbial populations separately for Bacteria and Archaea and for Fungi in both years for each location and land management using the Chao1 estimator of the number of SVs present in the sample, the Shannon entropy, the Simpson index of diversity and Faith’s phylogenetic diversity (Figure 2 and Supplementary Figure S2).

**Figure 2.**
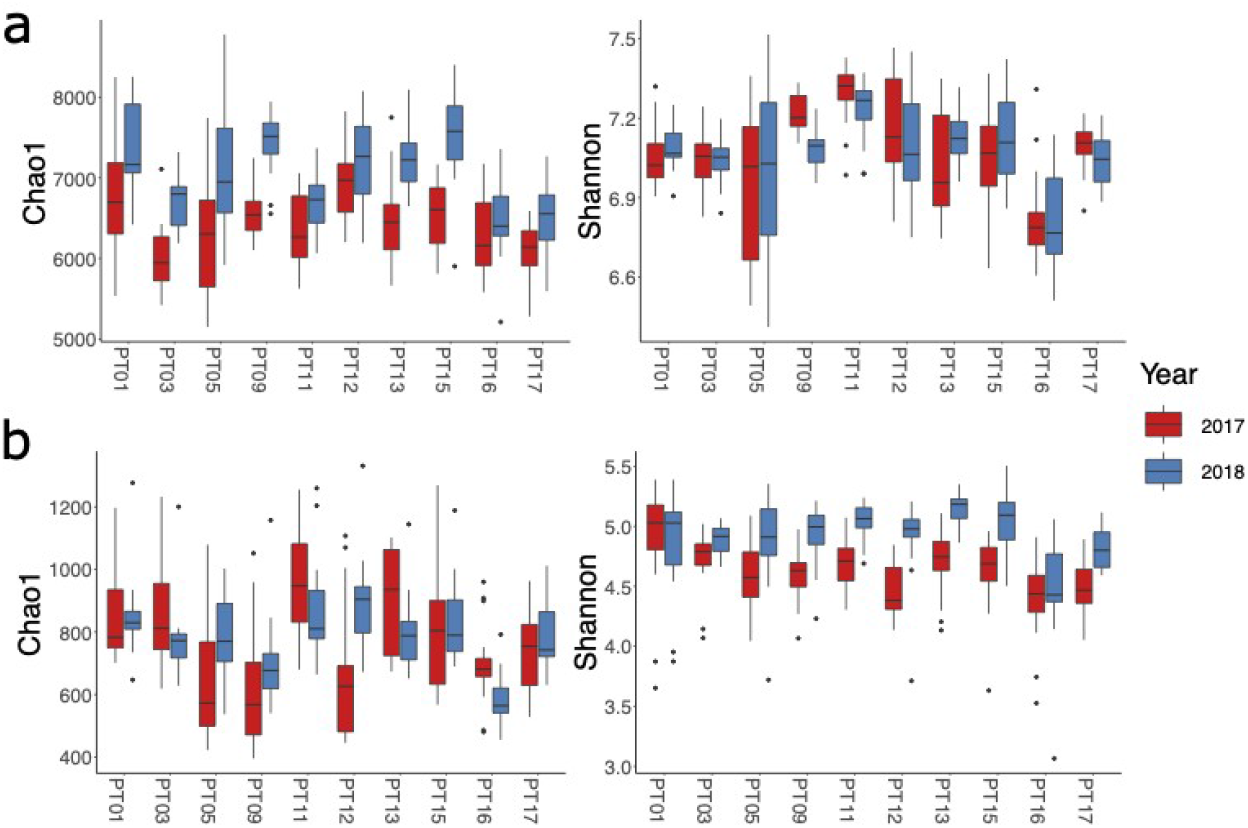
Richness and diversity of the bacterial a) and fungal b) components of the microbiota.

#### Bacteria and Archaea

Using a generalized linear model that accounts for differences between locations and land management (see Methods and Supplementary Materials), we found that the Chao1 estimator and Faith’s phylogenetic diversity were significantly higher in 2018 compared to 2017 (p-value < 2e-16 for both), while Simpson index was significantly lower in 2018 (p-value=9.26e-4) and there was no significant difference in Shannon entropy. These results were confirmed by a pair-wise comparison of the 168 sites for which we had samples both in 2017 and 2018 (p-value < 2.2e-16 for Chao1, < 2.2e-16 for Faith’s PD, and 2.47e-05 for Simpson index, respectively, one-tailed paired Wilcoxon rank-sum test). Again, the differences in the Shannon entropy between the two years were not significant.

The consistently higher value of the Chao1 estimator of species richness and of Faith’s phylogenetic diversity across all sites suggests that in each site there was a higher number of SVs that were present only in 2018 samples than of SVs that were present only in 2017 samples. To test this hypothesis, we computed, for each sampling site, the number of SVs that were specific to one of the two years and of those that were present in both (Figure 3a). We found that in most cases there was a higher number of SVs that were present only in 2018 than of SVs that were specific to 2017 (Supplementary Table 4). However, the year-specific SVs had a much lower relative abundance than SVs that were present in both years (Figure 3a), showing that the higher richness of 2018 samples was mainly due to rare taxa, i.e. to taxa that are present with a low number of individuals in each sample. Concerning the taxonomic distribution of year-specific and shared SVs (Figure 3b), we found that *Acidobacteria* and *Actinobacteria*, the two Phyla with the highest relative abundances (Figure 1a), had a higher number of taxa that were shared between the two years than of year-specific taxa, while the number of year-specific SVs was similar to the number of shared SVs for *Proteobacteria*, and higher than the number of shared SVs for *Planctomycetales* and *Bacteroidetes*. *Proteobacteria* was the phylum with the highest number of SVs that were present only in one of the two years in any given sampling site.

**Figure 3.**
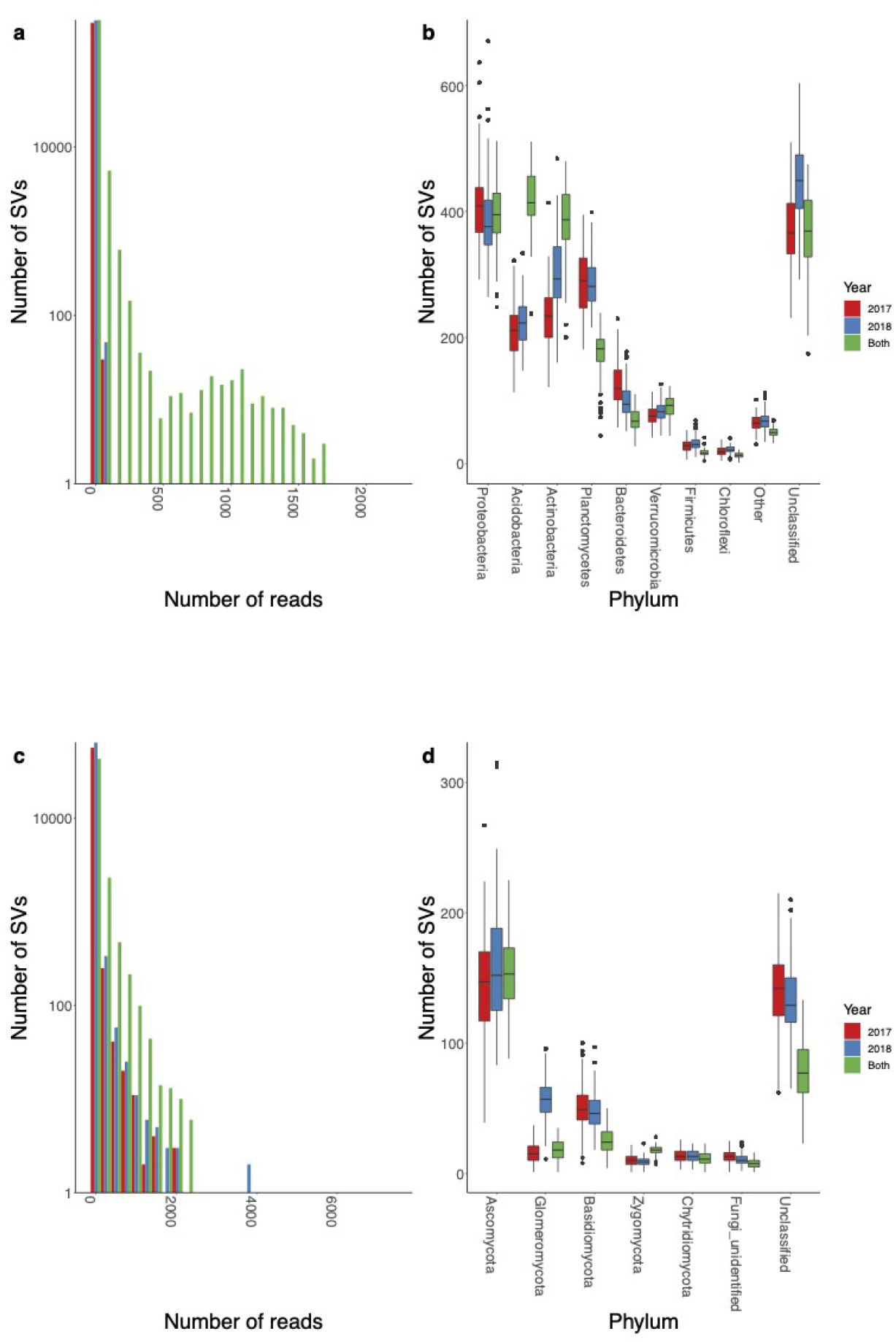
a) Number of bacterial Svs that, in a given site, were present only in one of the two years or in both, stratified by phylum. b) Number of bacterial Svs that, in a given site, were present only in one of the two years or in both, as a function of their abundance. c), d) Same as a), b), for fungi.

#### Fungi

As for Bacteria, we compared the alpha diversity of fungal communities using a generalized linear model that accounts for differences between locations and land management. We found that the Chao1 estimator of species richness was not statistically different between the two years while we found a significantly higher Faith’s phylogenetic diversity, although with high p-value (p-value=0.04924). These results were probably due to inconsistent variations across the different locations (Figure 2b). Indeed, while there were locations in which samples from 2018 had consistently higher Chao1 estimators for alpha diversity and Faith’s phylogenetic diversity across land management (*e.g.* PT12), in other cases the reverse was true, *i.e.* the Chao1 estimator of alpha diversity and Faith’s phylogenetic diversity of samples form 2018 were consistently lower that for 2017 across land management (*e.g.* PT16). On the other hand, we found that both the Shannon entropy and Simpson diversity index were higher in 2018 compared to 2017 (p-value <2e-16 and 2.06e-3, respectively). A pairwise comparison of the 161 sampling sites that, after rarefaction, had samples both in 2017 and 2018 confirmed that the increase of both Shannon entropy and Simpson diversity index was significant (p-value 3.7e-15 and 2.8e-08, respectively, one-tailed paired Wilcoxon rank-sum test).

Another striking difference between the fungal and bacterial component of the soil microbiome was evident by comparing the distribution of the abundances of the year-specific SVs (Figure 3c). In fungi, the SVs that were present only in one of the two years in each sampling site had a distribution of relative abundances that was similar to that of SVs that were conserved across years, including not only rare taxa, but also taxa with high relative abundances (Figure 3c), while in bacteria SVs present only in one of the sampling years included only SVs with a low relative abundance (Figure 3a). The taxonomic distribution of the year-specific and shared SVs (Figure 3d) showed that the Phylum *Ascomycota* had, in both years, a number of year-specific SVs that was comparable to the number of SVs that were present in both years, while the Phylum *Basidiomycota* was dominated by year-specific SVs. Interestingly, we found that in 2018 there was a general increase of the number of SVs from the Phylum *Glomeromycota* (Figure 3d). Indeed, the number of SVs from this Phylum that were specific to 2018 was higher both than the number of SVs specific to 2017 and the number of SVs that were present in both years.

### Soil microbiota maintain distinctive, site-specific characteristics across times

In order to characterize the relative importance of sampling site, location, land management and year of sampling to determine the differences between the samples, we computed the Bray-Curtis dissimilarity amongst all samples for Bacteria and Fungi. A PcoA analysis (Figure 4a) shows that while the different years induced some variability amongst the samples both for Bacteria and Fungi, samples still grouped according to location, suggesting that the variability between two consecutive years was comparable to the variability between the different samples from the same location and land management.

**Figure 4.**
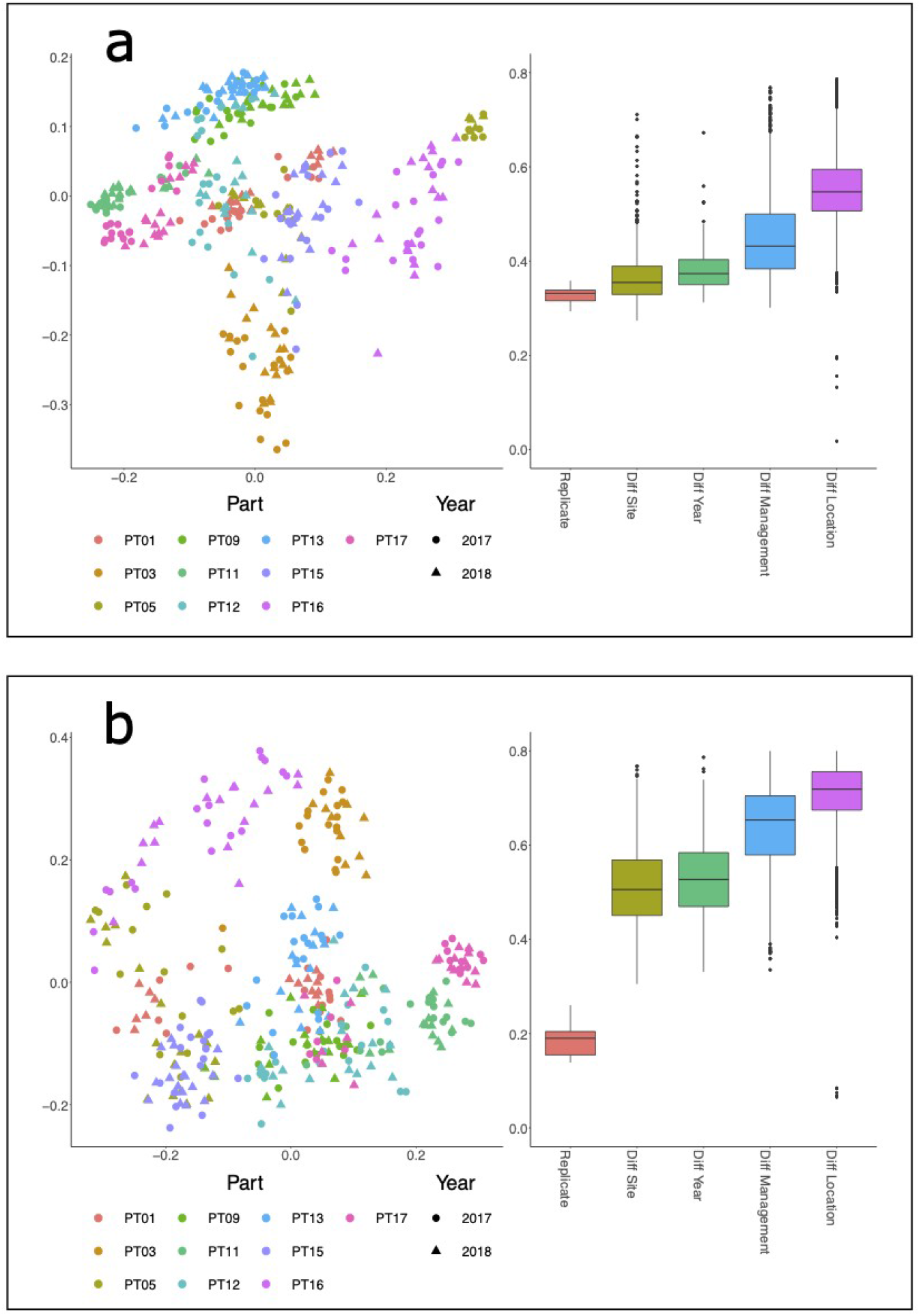
a) Left panel:PCoA of the Bray-Curtis dissimilarity for bacteria; Right panel: boxplots of the between-samples Bray-Curtis dissimilarities between pairs of samples for bacteria. In all comparisons only one factor is different, while all others are held constant. b) same as a), for fungi.

To quantify the relative importance of the different factors of the experimental design, namely sampling site, location, land management and year to determine the differentiation of the soil microbiota, we compared the distributions of the Bray-Curtis dissimilarities between pairs of samples where only one of the factors was varied, while all the other were held constant. The comparisons were between: i) samples from the same location, sampling site (and thus land management) and different year; ii) samples from the same location, year and different land management; iii) samples from the same land management, year, and different locations. These were compared with the dissimilarities between technical replicates. For both Bacteria and Fungi (Figure 5), the factors determining dissimilarity were, in growing order of importance, sampling site, year of sampling, land management and sampling location. All differences were statistically significant (Supplementary Table 4).

**Figure 5.**
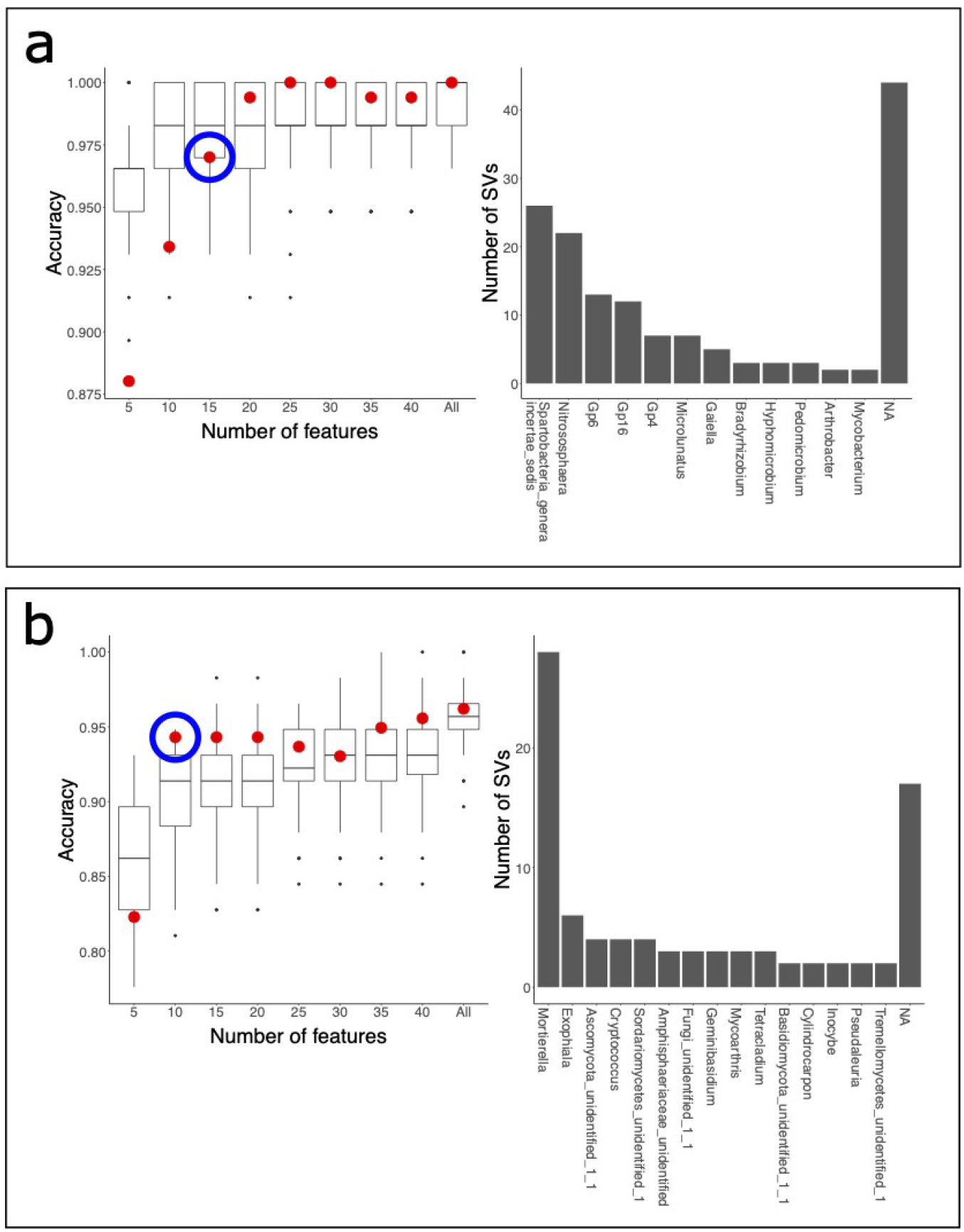
a) Left panel: Red dots are the prediction accuracy on the 2018 samples by a SVM trained over 2017 samples as a function of the number of selected bacterial SVs per site. The boxes are the cross-validation expected value of the accuracy of the SVM over the 2017 samples. The blue circle marks the minimal number of features that are needed for a 2018 prediction with an accuracy similar to the 2017 cross validation value. Right panel: Number of predictive SVs (corresponding to the blue dot in the left panel) classified at the Genus level. Only genera with at least two predictive SVs are shown. b) Same as a), for Fungi.

We used a Machine Learning approach to verify if the structure of the soil microbiota of each location maintained a set of organisms that distinguished it from the other locations and if these characteristic organisms were stable across different years. Specifically, we trained a set of classifiers based on Support Vector Machines that used the relative abundances of Bacteria and Fungi, respectively, to predict the location where a given sample had been collected. Using a cross-validation approach on 2017 data, we estimated that the mean expected Classification Accuracy (CA) was 0.989 (SD=0.0155; median 1.00) for Bacteria and 0.951 (SD=0.0267; median 0.948) for Fungi. We then trained one classification model for Bacteria and one for Fungi using all 2017 data, and used these models to predict location provenance of 2018 samples. We found that the CA of the 2018 samples was 1.0 for Bacteria and 0.96 for Fungi, showing that soil microbiota is highly associated with sampling location and that this association is stable over time.

In order to identify a reduced set of site-specific SVs that could be used to classify the samples according to their origin across different years, we used a three step procedure: i) first, we used the coefficients of the 2017 SVM models to rank the SVs in order of importance according to their contribution to the site classifier; ii) next, including the SV starting from the most relevant, we estimated how the CA depended on the number of included SVs using a cross-validation approach on the 2017 samples; iii) third, we used the same reduced set of SVs to predict the provenance of the 2018 samples using predictors trained on the 2017 samples. Using this procedure, we defined the set of SVs that characterize a given site as the minimal set for which the CA for the 2018 samples predicted using 2017 data exceeded the 25th percentile of the cross-validation expected CA. The 2017 expected CA (step ii), Figure 5) monotonically increased as a function of the number of included SVs to a mean value of 0.984 (median 0.983) for Bacteria and 0.93 (median 0.931) for Fungi using 40 SVs per site. The CA for 2018 samples was lower than the 2017 expected CA for the lower values of the number of included SVs, and exceeded the 25th percentile of the expected CA when more than 15 SVs per site were used for Bacteria, and 10 for Fungi. These were the minimal set of Bacterial or Fungal SVs for which the provenance of 2018 samples could be predicted using 2017 samples as training with performances that were similar to the cross validation expected value on a single year. Interestingly, the predictive SVs belonged to a limited number of taxa both for Bacteria and Fungi. Indeed, while 44/150 predictive bacterial SV could not be classified at the Genus level, the remaining 106 spanned only 13 Genera, including *Spartobacteria incertae sedis*, *Nitrososphaera*, *Gp6*, and *GP16*, which were represented by more than 10 SVs each. Of these, only SVs from *Spartobacteria incertae sedis* were discriminant for all sites. For Fungi, 28/100 predictive SVs belonged to the Genus *Mortierella*, which was also the only Genus that had predictive SVs for all locations.

## Discussion

In the last few years we have witnessed an increasing number of studies that use high throughput sequencing and culture-free approaches to study the composition of microbial communities in a variety of environments on a global scale. Thanks to this wealth of new data, we are starting to understand the major ecological drivers of microbial diversity and we have been able to greatly expand the census of known microbial species. The vast majority of the available data are from cross-sectional study designs, where environments are sampled across conditions at a given time. Much less experimental effort has been devoted to characterizing the stability of the microbiota over time, and how the variability due to sampling over different years time compares to other factors, like, in the case of soil, geography or land use. Soil bacterial communities have been shown to undergo significant shifts when sampling is repeated in different months within a year [15][35] In another study sampling contrasting seasons on a large spatial scale in wheat croplands across North China Plain, it was found that spatial variability was larger than temporal variability both for bacteria and fungi [16]. Inter-annual rates of community turnover have been shown to be much smaller than seasonal changes for fungi [36]. Here, we have sampled soil microbial communities in ten locations, in sites with three distinct land managements in each location, over two consecutive years.

Previously we found that the variability of fungal communities patterns were qualitatively different from what was found for bacteria. Despite the relatively limited geographical range sampled, these results were in striking agreement with the dominant role of a small number of taxa, in particular from the phylum Ascomycota [9]. Further, we found that geographical location, but not land use, had an impact on determining the size of the core soil mycobiome, indicating the importance of spatial processes in structuring the biogeographic pattern of soil fungal communities [17].

In this work we explored the effect of different years and studied how the year-to-year variability compares with other factors. We found that both for bacteria and fungi the differences due to different years were smaller than the differences due to land management or geographical location. These results show that, while vintage can cause significant shifts in grape microbiota even within small geographical scales [14], the soil microbiota of single vineyards is stable across consecutive years, probably due to the weaker influence of climate on bulk soil than on the more exposed grape surface. Comparing the species richness in the same site between two consecutive years, we found that there was a systematic difference in bacterial species richness and phylogenetic diversity that were consistently higher in 2018 samples regardless of location or land management, while for fungi we found that species richness and phylogenetic diversity changed in a site-dependent fashion. We found a general increase in Shannon entropy and Simpson diversity index for Fungi, indicating a decreasing role of the dominant species in the structure of the soil mycobiota in 2018 compared to 2017. Comparing meteorological data for the two years, we found that 2018 before the sampling date was characterized by a higher amount of accumulated rain compared to 2017. Few data exist regarding the relation between annual amount of rain and soil microbial richness and diversity, and most existing data report the effects of extreme phenomena, like drought or flooding [37]. In a global survey of the topsoil microbiota, Bahram et al. found that bacterial taxonomic diversity was negatively correlated with the mean annual amount of rain, while an inverse relation was found for fungi [3]. However, these data were obtained comparing samples from different locations, opening the possibility that these relations are due to general differences between wet and dry environments, and not to the effect of climatic variables alone. Here, by comparing two consecutive years in the same set of locations, we highlight the effects of yearly climate differences on soil microbiota. Our results might thus indicate a possible scenario in the case of short term climate change.

When we looked in detail at the composition of the yearly variable bacterial species, we found that these were mainly composed by rare taxa, *i.e.* taxa with low relative abundances, while the abundance distribution of yearly variable fungal species was similar to the abundance distribution of permanent species. Rare taxa are an integral component of microbial communities, that often display a long tail of low abundance species [38], and constitute a reservoir of microbial diversity that responds to environmental changes, thus contributing in an essential way to the dynamics of microbial communities [39]. Here, we found that a higher number of rare bacterial taxa were detected in 2018 compared to 2017, probably due to taxa that were below detection limit in samples from 2017, and that, due to different environmental conditions, grew in relative abundance in 2018. Interestingly, we found that the number of year-specific SVs was much lower than that of SVs that were conserved across years in *Acidobacteria* and *Actinobacteria*, the two Phyla with highest relative abundance, while was comparable or higher than that of SVs conserved across years for less abundant Phyla, like *Proteobacteria* and *Planctomycetales*.

For fungi, the situation was different. The fungal component of the microbiota had a simpler structure, with a smaller number of SVs than bacteria (compare, e.g., the values of the Chao1 estimator of species richness for Bacteria and Fungi, Figures 2a and 4a) and a higher spatial variability over short scales and temporal variability, as shown *e.g.* by the different scales in the distance distributions for bacteria and fungi (Figure 5), and the higher relative abundances of year-specific SVs for fungi compared to bacteria, (Figure 3). Thus, the role of stochastic fluctuations appears to be higher for Fungi than for Bacteria, with a more prominent role of temporal and spatial fluctuations, probably due to the lower complexity of soil fungal communities compared to bacterial communities.

A machine learning approach showed that a classifier trained over 2017 samples were able to correctly classify samples from the following year both using Bacterial and Fungal relative abundance data, showing that the soil microbiota has characteristics specific of the sampling location that are stable over consecutive years. Ranking SVs by their contribution to the classification, we found that predictive SVs belonged to a small number of Bacterial and Fungal genera, among which the most common were *Spartobacteria genera incertae sedis* and *Nitrososphaera* for Bacteria, and *Mortierella* for Fungi. *Spartobacteria* are a class of poorly characterized *Verrucomicrobia* that are highly abundant and ubiquitous in soil [40], particularly in grasslands where they appear to be well adapted to limited nutrient availability [41]. Species belonging to *Spartobacteria* have been found to be indicator species in acidoneutral Antarctic soils [42]. Abundance of *Spartobacteria* has been shown to increase with elevation in pasture soils in the Central European Alps [43]. Relative abundance of *Spartobacteria genera incertae sedis* in the rhizosphere has been found to positively correlate with plant growth in replanted apple orchards [44]. *Nitrososphaera* is a genus of *Archaea* that has been found at high relative abundance in vineyards soils and that contribute to nitrogen transformation [45]. *Mortierella* is a genus from Phylum *Zygomycota* that includes species of saprotrophs that live in soil. Members of this genus have been shown to be ubiquitous and present at high relative abundances in surveys of vineyard soils [46,47]. There are a few technical issues that might impact the results presented here. In particular, the presence of relic DNA has been shown to artificially inflate the estimated diversity (Carini et al. 2016). Although the effect of relic DNA on richness can act in either way depending on the dynamics of DNA degradation and has little effect on beta diversity [48], it is reasonable to expect that it can reduce the size of temporal fluctuations in soil microbial communities [49], thus dampening the year-to-year variability found in soil samples if the species abundance distribution is different between the two years [48]. For this reason, we expect that the systematic differences that we found between 2018 and 2017 samples are an underestimate of the real variations, and that further extension of the sampling on multiple years will help elucidate the impact of climate fluctuations on soil microbial communities.

It has been suggested that soil microbial communities are a reservoir of grape microbiota [14], and that vineyard soil microbiota could have an impact on wine fermentation [10]. It has also been shown that regional varieties of yeasts strongly contribute to wine regional characteristics [11], laying the base for the definition of a microbial terroir [50]. In order to contribute to the regional diversity of wine, the soil microbiota should be stable across different years, and the temporal variability should not be so large to wipe out the differences between different sites. We have shown that differences between consecutive years are smaller than those due to geographical factors even at short length scale, and that the structure of the soil microbiota is a signature of the geographical origin of the sample. Using a predictive model, we have shown that these site-specific features of the soil microbiota are stable across years, putting the concept of microbial terroir on a firmer ground.

## Materials and Methods

### Sample collection

The sampling sites were identified in 10 vineyards from 4 different locations (Ala, Besagno, Mori and S. Felice) because of their contiguity, at least along 20 meters, to perennial crop-covered surfaces. The experimental protocol set 3 sampling points respectively between the rows (V) and in the perennial crop area at a distance of 8 (P1) and 16 (P2) meters from the border of the vineyard [20]. Sampling was conducted in two consecutive years (2017 and 2018) in the same season (Supplementary Table 1). The dominant grass species in V sites were species belonging to the Poaceae family, while in P1 and P2 sites the dominant species were *Arrhenatherum elatius*, *Bromus erectus* and *Trisetum flavescens*. For each position 6 equally spaced sampling repetitions were performed, for a total of 180 samples for each sampling year.

Supplementary table 2 shows sites localization and technical characteristics of the vineyards (planting year, previous crop). All samples had a similar range of soil texture (loam, sandy clay loam, sandy loam and silty loam, see Supplementary Fig S1).

Quantity of soil organic matter (SOM), total nitrogen, total carbonate, and heavy metals (Cu and Zn) are reported in Supplementary Table 3. Samplings were executed collecting 20 cm of soil by means of a manual, one-piece, 7 cm diameter drill for loamy soils (Eijkelkamp, Edelman model). For chemical analysis and for taxonomic purposes of bulk soil the first 5 cm of soil were removed. Each sample consisted of 4 drillings that were homogenized in a signed plastic bag. From every one of them, a small volume of soil was collected in a 50 ml tube and chilled to 6/8°C during the sampling time after which they were frozen at −18°C.

### DNA extraction, library preparation and sequencing

The soil samples were freezed, dried and sieved with a 0.2 mm mesh size and stored at −80 °C until DNA extraction. Total DNA was extracted from 0.25 g of each composite soil sample using the PowerSoil DNA isolation kit (MO BIO Laboratories Inc., CA, USA) according to the manufacturer’s instructions. Total genomic DNA was amplified using primers specific to either the bacterial and archaeal 16S rRNA gene or the fungal ITS1 region. The specific bacterial primer set 515F (5’-GTGYCAGCMGCCGCGGTAA-3’) and the 806R (5’-GGACTACNVGGGTWTCTAAT-3’) was used [21] with degenerate bases suggested by [22] and [23]. Although no approach based on PCR amplification is free from bias, this primer pair has been shown to guarantee good coverage of known bacterial and archaeal taxa [24]. For the identification of fungi, the internal transcribed spacer 1 (ITS1) was amplified using the primer ITS1F (5’-CTTGGTCATTTAGAGGAAGTAA-3’) [19] and ITS2 (5’-GCTGCGTTCTTCATCGATGC-3’) [25]. All the primers included the specific overhang Illumina adapters for the amplicon library construction.

For the 16S V4 region each sample was amplified by PCR using 25 μl reaction with 1 μM of each primer. More in detail, 12.5 μl of 2x KAPA HiFi HotStart ReadyMix and 10 ul forward and reverse primers, were used in combination with 2,5 μl of template DNA (5-20 ng/μl). PCR reactions were executed by GeneAmp PCR System 9700 (Thermo Fisher Scientific) and the following cycling conditions: initial denaturation step at 95 °C for 5 minutes (one cycle); 28 cycles at 95 °C for 30 seconds, 55 °C for 30 seconds, 72 °C for 30 seconds; final extension step at 72 °C for 5 minutes (1 cycle).

For the ITS1 region each sample was amplified by PCR using 25ul reaction with 10 μM of each primer. More in detail 22 μl of premix FastStart High Fidelity PCR System (Roche) and 2μl forward and reverse primers, were used in combination with 1 μl of template DNA (5-20 ng/μl). PCR reactions were executed by GeneAmp PCR System 9700 (Thermo Fisher Scientific) and the following cycling conditions: initial denaturation step at 95 °C for 3 minutes (one cycle); 30 cycles at 95 °C for 20 seconds, 50 °C for 45 seconds, 72 °C for 90 seconds; final extension step at 72 °C for 10 minutes (1 cycle).

The amplification products were checked on 1.5 % agarose gel and purified using the Agencourt AMPure XP system (Beckman Coulter, Brea, CA, USA), following the manufacturer’s instructions. Afterward, a second PCR was used to apply dual indices and Illumina sequencing adapters Nextera XT Index Primer (Illumina), by 7 cycles PCR (16S Metagenomic Sequencing Library Preparation, Illumina). The amplicon libraries were purified using Agencourtusing the Agencourt AMPure XP system (Beckman), and the quality control was performed on a Tapestation 2200 platform (Agilent Technologies, Santa Clara, CA, USA). Finally, all barcoded libraries were pooled in an equimolar way and sequenced on an Illumina® MiSeq (PE300) platform (MiSeq Control Software 2.5.0.5 and Real-Time Analysis software 1.18.54.0).

### Bioinformatic processing of the sequences

The sequences were assigned to samples using sample-specific barcodes and saved in FASTQ-formatted files. Sequences were deposited to the European Nucleotide Archive (ENA) with study accession PRJEB31356. Raw data FASTQ files were analyzed using the software pipeline MICCA [26] v. 1.7.2.

Raw overlapping 16S paired-end reads were assembled (merged) using the procedure described in [27]. Paired-end reads with an overlap length smaller than 200 bp and with more than 50 mismatches were discarded. After trimming forward and reverse primers, merged reads shorter than 250 bp and with an expected error rate higher than 0.5% were removed.

Filtered sequences were clustered into sequence variants (SVs) using the UNOISE3 denoising algorithm available in MICCA. OTUs were taxonomically classified using the Ribosomal Database Project (RDP) Classifier [28] v2.11. Multiple sequence alignment (MSA) was performed on the denoised reads applying the Nearest Alignment Space Termination (NAST) [26,29] algorithm and the phylogenetic tree was inferred using FastTree [30] (v2.1.8).

Raw overlapping ITS paired-end reads were merged and merged sequences with an overlap length smaller than 100 bp and with more than 32 mismatches were discarded. After primers trimming, merged reads shorter than 150 bp and with an expected error rate higher than 0.5% were removed. Filtered sequences were clustered into SVs using the UNOISE3 denoising algorithm and SVs were taxonomically classified using the RDP Classifier v2.11 and the UNITE [31] database. To compensate for different sequencing depths, samples were rarefied to an even depth of 15,000 reads for both 16S and ITS sequences. Samples with less than the minimum number of reads were discarded. Prior to rarefaction, sequencing reads from control runs (see section on Technical replicates) were merged with actual runs for those samples that were sequenced multiple times. Since P2 sampling sites of neighbouring PT05 and PT15 sites coincide, a single sequencing run was used for both PT05 and PT15 sites in diversity and distance calculations. These samples were not considered in the training and validation of the SVM classifiers.

### Technical replicates

Soil samples from 2017 and 2018 were sequenced in different times in several sequencing runs. To exclude that batch effects could affect our estimates of the differences between the richness and composition of the soil microbiota in the two different years, and of the relative importance of time, sampling site, and land use, we resequenced 10 samples from 2017 and 10 from 2018, one for each location, in a single sequencing run (hereinafter, for brevity, “control run”), and compared alpha and beta diversity indexes between these control samples and the corresponding actual samples. We found (see Supplementary Material) that there was a good degree of correlation between observed number of SVs, Chao1 estimator of species richness, Shannon entropy and Simpson diversity index in control and actual samples for Bacteria, and, to a lower degree due to higher sensitivity to rarefaction, for fungi. For beta diversity, the Bray Curtis dissimilarities between pairs of actual samples were always highly correlated with the dissimilarities between corresponding pairs of control samples, with minimal variance introduced by rarefaction.

### Statistical analysis of the data

Biom files and rooted phylogenetic trees (used in the calculation of beta diversity) were imported into R v4.0.0 using the *phyloseq* package [32] v1.32.0 for downstream statistical analysis. Alpha diversity was calculated using the Chao1 estimator [33] and the Shannon entropy [34]. Generalized linear models for alpha diversity were determined using the *glm* function in R. Contrasts were calculated using the package *emmeans* v1.5.3. Beta-diversity was calculated using the Bray Curtis distance.

Linear kernel Support Vector Machines encoded the LiblineaR v2.10-8 software package were trained using the R package mlr3 v0.8.0 through the mlr3extralearners v0.1.0 interface. Classification Accuracy was estimated by 50 folds cross validation by random splitting the dataset in 2/3 training and 1/3 test set. Tuning of the C parameter of the SVM was accomplished in each iteration using an inner 3-fold cross validation loop. Parameter search space was C=0.01,0.1,1,10,100,1000.

### Meteorological data

Meteorological data from the sampling areas were recorded daily in four stations located in close proximity of the vineyards (see Supplementary Informations). Based on location, the association between meteorological stations and sampling locations was as follows: Station 1 - Part05, Part09, Part12, Part15; Station2 - Part03, Part11, Part16, Part17; Station 3 - Part13; Station 4 - Part01.

## Availability of data and material

Raw sequencing data along with geographical and physico-chemical information are available at the European Nucleotide Archive (https://www.ebi.ac.uk/ena) under the study id PRJEB31356. Meteorological data are available as supplementary material.

## Acknowledgements

The meteorological data were kindly provided by “Unità Agrometeorologia e Sistemi Informatici” of the Edmund Mach Foundation (https://meteo.fmach.it/); furthermore authors would also thanks the “Consorzio vini del Trentino” (https://www.vinideltrentino.com/) for the sampling sites availability.

## Authors’ contributions

AC: Conceptualization. Resources. Writing - original draft. Writing - review and editing. Data curation.
EC: Conceptualization. Resources. Writing – review and editing.
DA: Formal analysis. Data curation. Software. Writing – review and editing.
ES: Investigation.
MP: Investigation. Resources.
CI: Writing – review and editing.
RZ: Investigation. Writing – review and editing.
CD: Conceptualization. Resources. Supervision. Formal analysis. Writing – original draft. Writing – review and editing.

